# Double Imprinted Nanoparticles for Sequential Membrane-to-Nuclear Drug Delivery

**DOI:** 10.1101/2023.07.19.549711

**Authors:** Pankaj Singla, Thomas Broughton, Mark V. Sullivan, Saweta Garg, Rolando Berlinguer-Palmini, Priyanka Gupta, Francesco Canfarotta, Nicholas W. Turner, Eirini Velliou, Shoba Amarnath, Marloes Peeters

## Abstract

Nanoparticles functionalized with specific receptors (*e.g.,* antibodies, peptides) are used for targeted drug delivery of anti-cancer agents but their side effects include hypersensitivity reactions, toxicity, inflammation, and life-threatening allergic reactions (Anaphylaxis) [1,2]. Consequently, double imprinted molecularly imprinted nanoparticles (nanoMIPs) against a linear epitope of breast cancer cell receptor estrogen alpha (ERα) and loaded with an anti-cancer agent (doxorubicin, DOX) are synthesized via a solid-phase approach. Surface plasmon resonance (SPR) measurements reveal that the produced nanoMIPs exhibit K_D_ values of 19 nM (against the epitope used for imprinting) and 10 nM (ERα receptor), and thus rival the affinity of nanoparticles decorated with natural affinity reagents (*e.g.,* antibodies, peptides), whilst offering the advantages of low-cost and enhanced cellular uptake due to the receptor mediated endocytosis. We present the results of *in vitro* flow cytometry that DOX loaded nanoMIPs can preferentially bind to MCF-7 (ERα positive) breast cancer (BC) cells vs MDA-MB-231 (ERα negative) BC cells. Confocal imaging witnessed the above results and showed the sequential movement of the DOX loaded nanoMIPs from membrane to the nucleus of MCF-7 BC cells and achieve delivery of DOX once internalised in the cells (directly to the nucleus). As a result, enhanced cell toxicity in MCF-7 cells (∼80%) as compared to MDA-MB-231 cells (∼15%) is observed via MTT (3-(4,5-dimethylthiazol-2-yl)-2,5-diphenyl-2H-tetrazolium bromide) cytotoxicity assay in a time dependent manner. Overall, this study provides a promising approach for the targeted drug delivery of chemotherapeutic drugs to breast cancer cells, which has the potential to significantly improve patient outcome whilst also reducing debilitating side effects of current treatment.

## 1.0 Introduction

Breast cancer (BC) is the most frequently diagnosed cancer worldwide; with 2.3 million new cases in 2020, this accounts for 1 in 8 cancer diagnoses [3]. It is a highly heterogeneous disease that can be caused by a variety of distinct genetic alterations in mammary epithelial cells, requiring a combinatorial evaluation of the histopathology of the primary tumour and of the expression pattern of hormone receptors to determine the optimal patient treatment plan. The majority (70%) of BCs are estrogen receptor positive (ER+), meaning the cancer is fuelled by the estrogen hormone [3,4]. ERα is a central transcription factor that is often overexpressed in BC, but also in ovarian, endometrial and prostate cancers [5], and plays a crucial role in breast tumorigenesis and proliferation of BC cells.

Traditionally, ERs have been considered nuclear receptors, but it has been well-documented that these receptors are also present on the membrane and in the cytoplasm. Nuclear ERα stimulates gene expression changes that promote cell cycle progression [6,7,8]. Moreover, the binding of ligands to plasma membrane ERα triggers rapid cellular changes through second messenger pathways, which also contribute to transcriptional effects of estrogen by regulating processes such as proliferation, cell migration, and development [9,10]. Membrane-bound ERα receptors are associated with caveolae that help in their internalization through dynamin-mediated processes and play a crucial role in initiating ERα signalling [11] Endocrine therapy, consisting of for instance tamoxifen and fulvestrant, is the preferred first line of treatment for ER-positive BC [12]. Chemotherapeutic drugs may also be prescribed alone or in combination with endocrine therapy [13]. Additionally, chemotherapy may be used after surgical resection or in the case of metastatic BC, with doxorubicin (DOX) being one of the most frequently prescribed chemotherapy drugs for solid breast tumours [14]. However, all current treatment options come with significant challenges and side effects, such as drug resistance, severe toxicity (for example, neurotoxicity and cardiac toxicity including irreversible heart injury) and allergic reactions (fainting, sweating) resulting from chemotherapy’s non-selective behaviour [14,15,16].

Nanoparticles are currently being utilized in the field of nanomedicine to enable targeted drug delivery of chemotherapeutic drugs, leading to more effective treatment of cancer while minimizing harmful side effects [17]. To achieve targeted drug delivery, nanoparticles (e.g gold nanoparticles, liposomes) are functionalized with targeting agents which can be broadly classified as proteins (antibodies and their fragments), nucleic acids (aptamers), or other receptor ligands (peptides, vitamins, and carbohydrates) [18,19]. Enhertu is a prime example of a commercial antibody drug conjugate for BC treatment: this product composed of trastuzumab antibodies conjugated with deruxtecan and are designed to specifically target and treat metastatic HER2 (human epidermal growth factor receptor 2)-positive BC. Another example is Nab-paclitaxel (Abraxane), FDA approved albumin-bound nanoparticles for the delivery of paclitaxel used as second-line treatment for metastatic BC [20,21]. However, these formulations all rely on biological counterparts, which may encounter significant limitations such as high cost (minimum ∼$100,000 per treatment for Enhertu) limited *in vivo* stability, inherent heterogeneity of biologics which can lead immune intolerance, and inability to recognize altered peptide antigens. These drawbacks are overcome using molecular imprinted polymeric nanoparticles (nanoMIPs), which are highly selective, cost-effective, and robust, and can serve as an alternative for targeted drug delivery. These synthetic receptors are small porous polymeric nanostructures containing specific binding sites for a particular target [22,23,24]. These materials possess several advantages such as excellent chemical and biological stability, biocompatibility, versatility, and fast binding kinetics [25,26,27]. Solid phase synthesis for the manufacturing of nanoMIPs has led to the production of high affinity materials with homogeneous binding distribution and excellent biocompatibility due to the use of the solid-phase as an affinity medium [28].

There are some examples of hybrid core-shell systems where drug-loaded nanoparticles are coated with MIPs to facilitate targeted drug delivery to cell-specific receptors [29, 30]. However, a more scalable and straightforward approach would be to design and synthesize double imprinted nanoMIPs. In this work, we have developed DOX loaded nanoMIPs for its targeted delivery to ERα positive BC cell lines to improve breast cancer treatment via double imprinting with an epitope of ERα in the presence of DOX. Via this innovative synthesis approach, we have streamlined the manufacturing to a one-stage process, which will significantly lower production cost and development time. There is one report on double imprinted nanoparticles where drug delivery of DOX is explored via targeting the membrane receptor epidermal growth factor receptor (EGFR). As this material only bound to the receptor and was not internalised by the cells, the therapeutic effect in this case can be limited [31]. Our work introduces a novel approach by successfully targeting the membrane receptor ERα and facilitating the translocation of nanoMIPs towards the nucleus, which paves the for the selective drug delivery of traditionally “undruggable” targets with MIPs.

In this context, nanoMIPs have been developed to target ERα and were tested on two different BC cell lines: MCF-7 (ERα positive) and MDA-MB-231 (ERα negative), which represent different subtypes of BC and thus require different treatment in clinical practice. The study showed that nanoMIPs designed for ERα successfully bound to this receptor on the membrane and were subsequently internalized, thus facilitating highly specific intracellular delivery of DOX. By enabling nuclear delivery of DOX, the nanoMIPs elicited significantly higher cytotoxicity (80%) towards the MCF-7 cell line as compared to the negative control cell line MDA-MB-231 (15%).

Furthermore, the efficacy was evaluated in 3D scaffolds of BC cell lines, which provide a better representation of BC progression and response to drugs *in vivo.* 3D scaffolds provide a remarkable platform for in vitro testing of these nanoparticle formulations by effectively mimicking the complex tumour microenvironment found in living organisms. Thus, these engineered nanoMIPs have great potential as a targeted drug delivery vehicle in the field of precision medicine and can easily be expanded to other cancer types given the versatility of the approach used to manufacture these engineered nanoparticles.

## 2.0 Results and discussions

### 2.1 Solid phase synthesis and characterization of nanoMIPs

We have produced nanoMIPs to target the human ERα receptor protein (595 amino acids) with a molecular weight of 66 kDa (ERα66 wild type). A linear epitope sequence SHSLQKYYITGEAEGFPATV (576-595 amino acids at the C-terminus, corresponding to the binding region of Anti-ERα Antibody F-10, sc-800, Santa Cruz Biotechnology, Inc. USA) on helix 12 of human ERα receptor (UniProtKB-P03372) was selected [32]. A cysteine residue was added to the N-terminus of this peptide sequence for binding to the solid support used to produce the nanoMIPs, and its attachment was confirmed via a BCA assay (**Figure 1a**). To circumvent the limitations connected with the use of whole proteins, the epitope imprinting approach offers several advantages *viz.* lower costs, compatibility with different synthetic conditions (pH, temperatures, and solvents), selectivity towards target, greater versatility, no need for costly and lengthy protein purification steps [33,34]. The epitope-imprinting method is well-established and MIP-based sensors produced with this approach have successfully achieved full protein identification [24,35,36].

**Figure 1:**
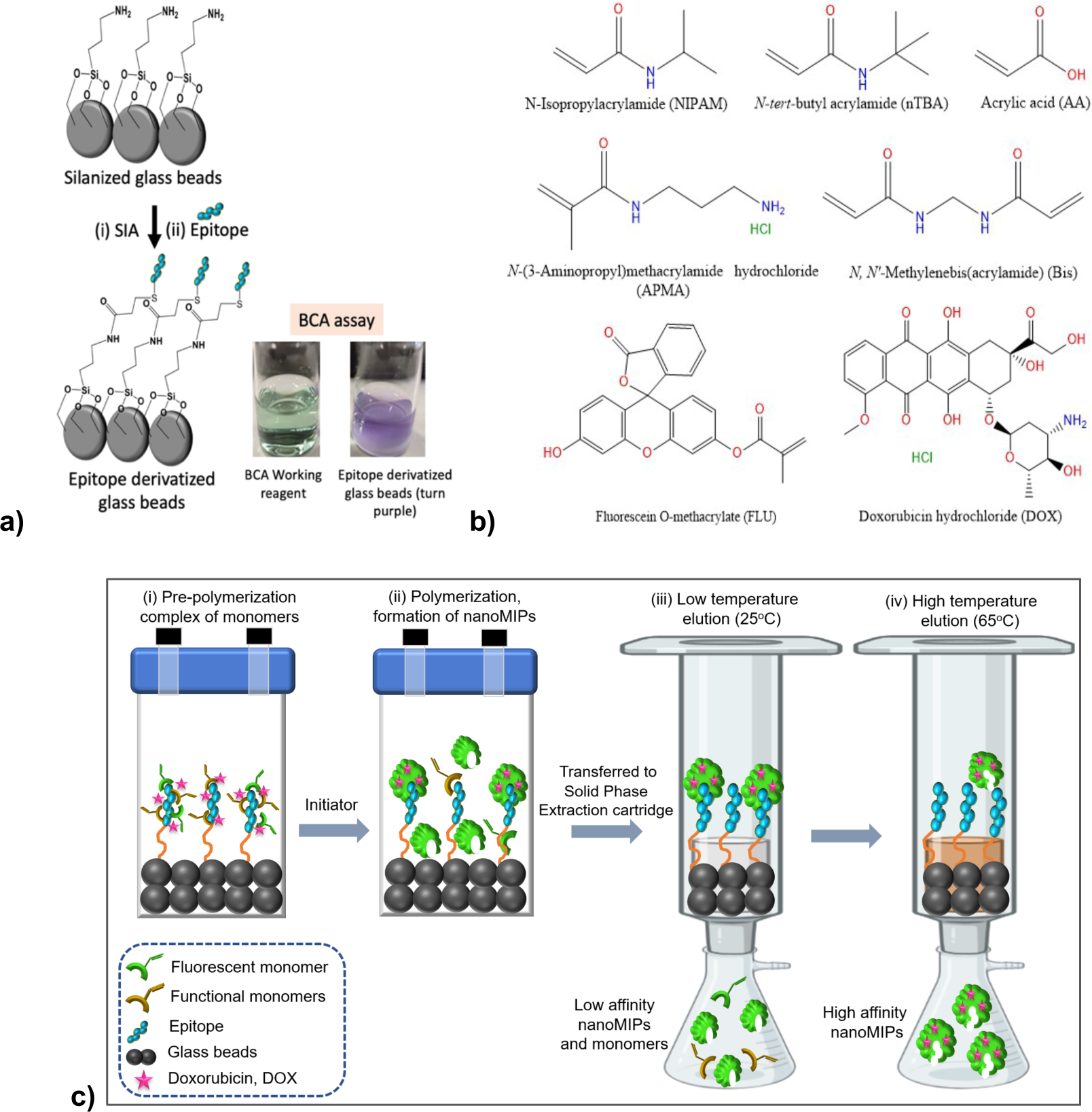
**a)** Molecular structures of different functional monomers, fluorescent monomer (Fluorescein-o-methacrylate), and DOX **b)** Schematic of conjugation of cysteine modified epitope with silanized glass beads and confirmation through bicinchoninic acid (BCA) assay (green to purple in the presence of epitope modified glass beads) **c)** Schematic of synthesis of double imprinted nanoMIPs: the primary template is the epitope conjugated on silanized beads, whilst DOX acts as secondary template in solution.

A variety of monomers were employed, comprising *N*-Isopropylacrylamide (NIPAM) for hydrogen bonding, *N*-*tert*-Butylacrylamide (nTBA) for hydrophobic interactions, and *N-*(3-Aminopropyl)methacrylamide hydrochloride (APMA), and acrylic acid (AA) for ionic interactions. Fluorescein-o-methacrylate was added in the monomer mixture to obtain fluorescent active nanoMIPs that enable tracking of the nanoparticles in the 2D cell lines and 3D scaffolds when imaging of the system. The molecular structures of the monomers and DOX are depicted in **Figure 1b**, while the composition of various batches of nanoMIPs and NIPs produced in this study is presented in **Table S1**. Fluorescein tagged nanoMIPs without DOX (unloaded) are referred to as FLU-nanoMIPs, whereas DOX loaded, and fluorescein tagged DOX loaded double imprinted nanoparticles are named DOX-nanoMIPs and FLU-DOX-nanoMIPs respectively. Non-imprinted nanoparticles (NIPs, named as FLU-NIPs and FLU-DOX-NIPs were also prepared through solid phase synthesis approach using silanized beads and serve as a reference. The solid-phase synthesis method for the manufacturing of the different nanoMIPs is depicted in **Figure 1c**.

The hydrodynamic diameter (*D_h_*) of FLU-nanoMIPs, DOX-nanoMIPs and FLU-DOX-nanoMIPs were found to be 121 ± 3 nm (PDI=0.115), 141 ± 3 nm (PDI=0.118) and 168 ± 2 nm (PDI=0.127) respectively (**Figure 2a**). These results showed that the loading of DOX into nanoMIPs increased *D_h_* of nanoMIPs, due to accommodating fluorescein-o-methacrylate and DOX within the nanoMIPs. PDI values of these nanoMIPs were found to be less than 0.2 suggesting that these nanoparticles are homogeneous.

**Figure 2:**
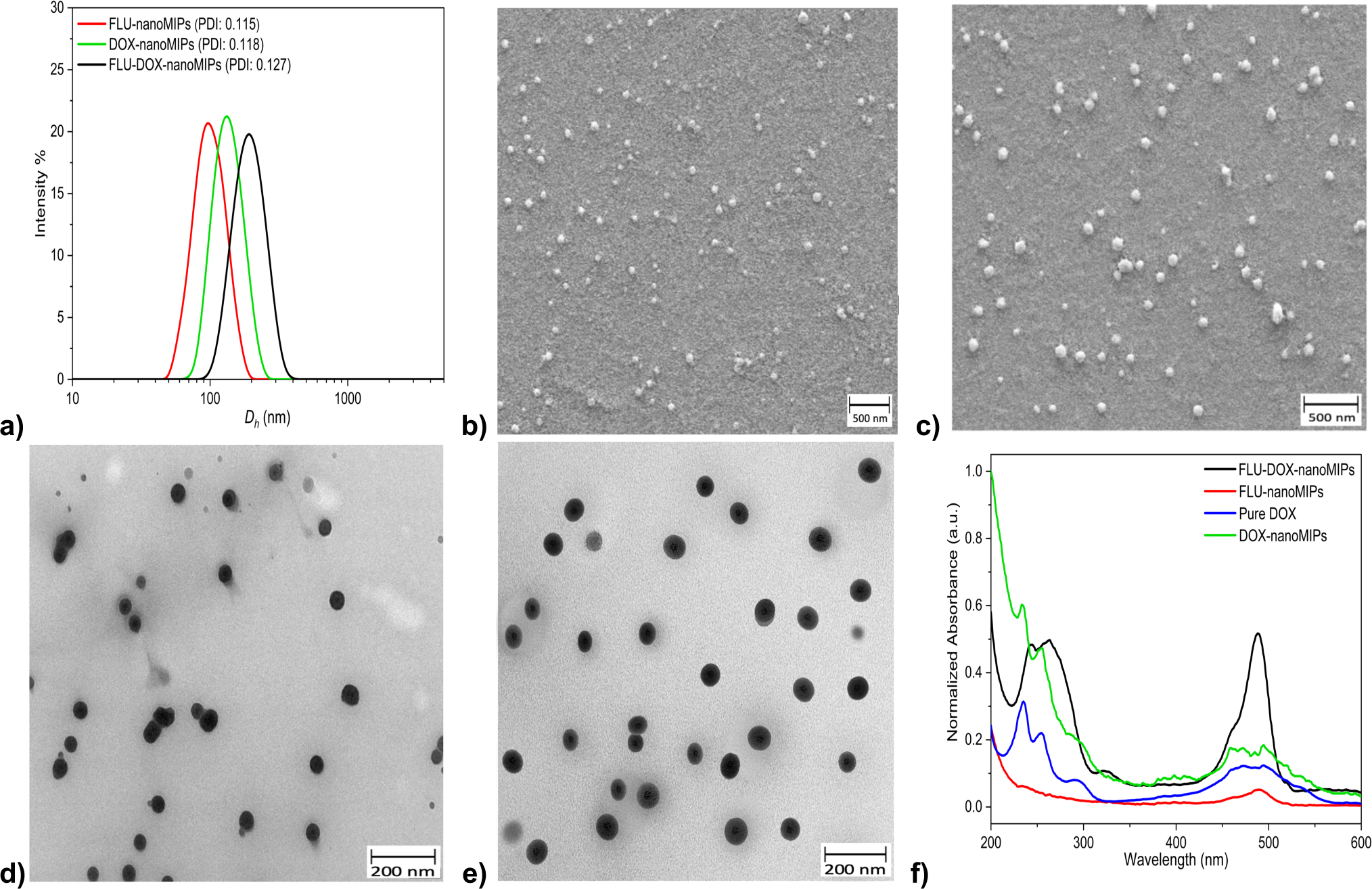
Characterization of different nanoMIPs (**a**) Intensity weighted size distribution plot (DLS measurement) FLU-nanoMIPs (*Dh* = 121.16 ± 2.70), DOX-nanoMIPs (*Dh* = 141 ± 2.76) and FLU-DOX-nanoMIPs (*Dh* = 168 ± 2.24) at 25^◦^C. The Z-average *Dh* of the nanoMIPs was determined using cumulant analysis by the equipment, and the standard deviation was calculated based on 5 measurements; representative SEM images of **b)** FLU-nanoMIPs **c)** FLU-DOX-nanoMIPs; TEM image (25000x) of **d)** FLU nanoMIPs, **e)** FLU-DOX-nanoMIPs; **f)** UV-spectra of Pure DOX, FLU-nanoMIPs, DOX-nanoMIPs and FLU-DOX-nanoMIPs.

The average size from the scanning electron microscopy (SEM) measurements for the FLU-nanoMIPs and FLU-DOX-nanoMIPs were observed to be 46±11 nm and 60±10 nm (**Figure 2b, c**) respectively, whereas DOX-nanoMIPs showed the size of 58±6 **(Figure S1 a)**. This was in line with the transmission electron microscopy (TEM) results, where the size of FLU-nanoMIPs and FLU-DOX-nanoMIPs was found to 40±6 nm and 58±6 respectively (**Figure 2 d, e**) and DOX-nanoMIPs showed size of 57±0.9nm **(Figure S1 b)**. Moreover, both methodologies confirmed that the nanoMIPs exhibited a spherical morphology. The larger size obtained by DLS measurements can be contributed to the hydration of the nanoMIPs as the AA and NIPAM monomers significantly in the liquid state. On the contrary, the size obtained by SEM and TEM is indicative of the nanoMIPs size in the dry state [37]. The variation in sizes observed between nanoMIPs and DOX-loaded nanoMIPs was primarily attributed to the incorporation of DOX into the nanoMIPs, resulting in an overall increase in their size. UV-spectres of different batches DOX loaded nanoMIPs have been shown in **Figure 1 f** and estimation of DOX loading was performed by determining the amount of imprinted drug using a calibration curve (**Figure S2**), as well as assessing the loading efficiency and loading capacity (**Table-S2**). The loading of DOX into DOX-nanoMIPs, FLU-DOX-nanoMIPs, and FLU-DOX-NIPs was found to be 17.28 ± 0.1, 19.33 ± 0.16, and 18.37 ± 0.12 μg/100 μg of nanoMIPs, respectively, with corresponding drug loading efficiencies of 57.6 ± 0.33 %, 64.43 ± 0.53 %, and 61.12 ± 0.40 %. This DOX loading is comparable to the Doxosome™ (DOX 20 μg/ ∼160 μg of lipids), which is commercially available liposomal nano formulation specifically designed for research and development purposes [38].

### 2.2 Binding performance and selectivity of nanoMIPs

The prepared nanoMIPs were tested for their binding affinity towards the ERα protein (68kDa) and the ERα template epitope used in the imprinting process via surface plasmon resonance (SPR) in a solution of PBS pH 7.4 and 0.01% Tween 20. To compare the effect of DOX loading on nanoMIPs, two different types of nanoMIPs (FLU-nanoMIPs and DOX-FLU-nanoMIPs) were explored for the affinity against both ERα protein and ERα epitope (CSHSLQKYYITGEAEGFPATV). The nanoMIPs were immobilized onto carboxymethyl dextran hydrogel-coated gold (Au) chips through EDC-NHS chemistry, where the amine moiety of nanoMIPs was crosslinked with the carboxymethyl dextran on the Au chip [39]. The sensograms depicted in **Figure 3a**, correspond to the association and dissociation between the ERα protein to FLU-nanoMIPs, with a similar trend observed for the other set of experiments for FLU-nanoMIPs with template epitope (**Figure 3b**), FLU FLU-DOX-nanoMIPs with ERα protein and template epitope (**Figure 3c and d**) respectively. The increase in signal referred to the association of epitope/ERα protein to the nanoMIPs and the association constant (*K_on_*) have been determined. The decrease in the signal showed the dissociation phase by which the dissociation constant was calculated (*K_off_*). The binding affinity of a molecular interaction was quantified by the equilibrium dissociation constant (*K_D_*) calculated from the values of *K_off_ / K_on_*. Response increases linearly in case of template epitope/ ERα protein (from concentration range of 4 to 64 nM) for both FLU-nanoMIPs and FLU-DOX-nanoMIPs. The *K_D_* values for the entire ERα protein was found to be 14.7 nM and 10.8 nM for FLU-nanoMIPs and FLU-DOX-nanoMIPs, respectively. In case of template epitope, *K_D_* values came out to be 19.2 nM and 16.2 nM for FLU-nanoMIPs and FLU-DOX-nanoMIPs respectively.

**Figure 3:**
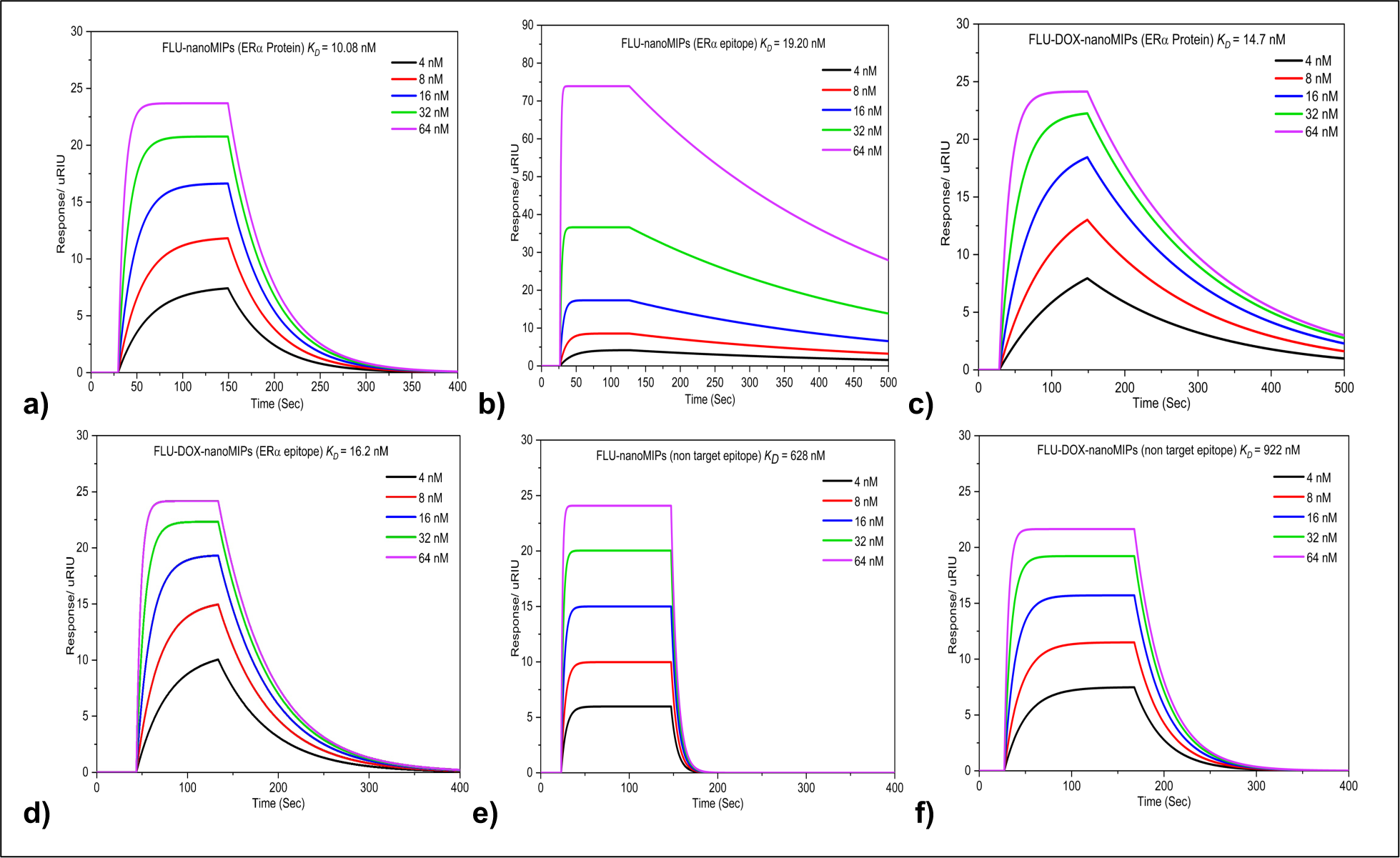
Typical sensorgrams depicting binding of different nanoMIPs **a)** FLU-nanoMIPs with ERα protein **b)** FLU-nanoMIPs with ERα epitope **c)** FLU-DOX-nanoMIPs with ERα protein **d)** FLU-DOX-nanoMIPs with ERα epitope **e)** FLU-nanoMIPs with non-target epitope, and **f)** FLU-DOX-nanoMIPs with non-target epitope.

Typically, antibodies have *K_D_* values falling within the low micromolar range to nanomolar range, indicating moderate to high affinity for the antigens they bind to. However, antibodies that exhibit even stronger binding affinity are considered to be high-affinity antibodies, with *K_D_* values falling within the low nanomolar to sub-nanomolar range [40]. Our results revealed that the binding affinity of our nanoMIPs was comparable to that of antibodies. Selectivity is a crucial factor in the development of nanoMIPs as a targeted drug delivery system because it determines the ability of nanoMIPs to selectively bind and deliver drugs to the desired target, such as cancer cells, while minimizing their toxicity to healthy tissue [41]. The selectivity of nanoMIPs was evaluated by using the nontarget epitope SSERIDKQIRYILDGISALR (epitope of interleukin 6), which has a similar molecular weight (M_w_) of 2333.64 Da. The results showed that the *K_D_* values for the nontarget epitope were approximately 62 times higher for both FLU-nanoMIPs and FLU-DOX-nanoMIPs as compared to target ERα protein (**Figure 3e and f**). Specifically, results found that a 62-fold higher concentration of the nontarget is required to occupy 50% of the nanoMIPs. These results demonstrated the high selectivity of the nanoMIPs for the target ERα protein, which is critical for their effectiveness as a targeted drug delivery system.

### 2.3 *In-vitro* cell binding and specificity of nanoMIPs

Two different BC cell lines MCF-7 (ERα positive) and MDA-MB-231 (ERα negative) have been chosen to visualize the specific binding of nanoMIPs using flow cytometry. Flow cytometry binding for FLU-nanoMIPs, FLU-DOX-nanoMIPs and DOX-nanoMIPs at 10μg/mL concentration with MCF-7 and MDA-MB-231 are represented in **Figure 4 a, b and c** respectively. These results demonstrate that there is significantly higher binding of the nanoMIPs (10μg/mL) to MCF-7 cells in comparison to MDA-MB-231 cells. The higher the fluorescence intensity, the higher is the binding of nanoMIPs towards the cells overexpressing ERα. The mean fluorescence intensity (MFI) plots of the FLU-nanoMIPs (P ≤ 0.001), the FLU-DOX-nanoMIPs (P ≤ 0.0001), and the DOX-nanoMIPs (P ≤ 0.01) shown in **Figure 4 d-f** demonstrated that MCF-7 cells exhibited ∼4 folds greater number of MFI positive cells than MDA-MB-231 cells. However, low levels of binding with MDA-MB-231 cells have also been encountered, which can occur since some report suggest that there is small amount of ERα present in that cell line [42–43], in addition to the fact that all nanoparticles have some unavoidable non-specific binding to the surface of cells.

**Figure 4:**
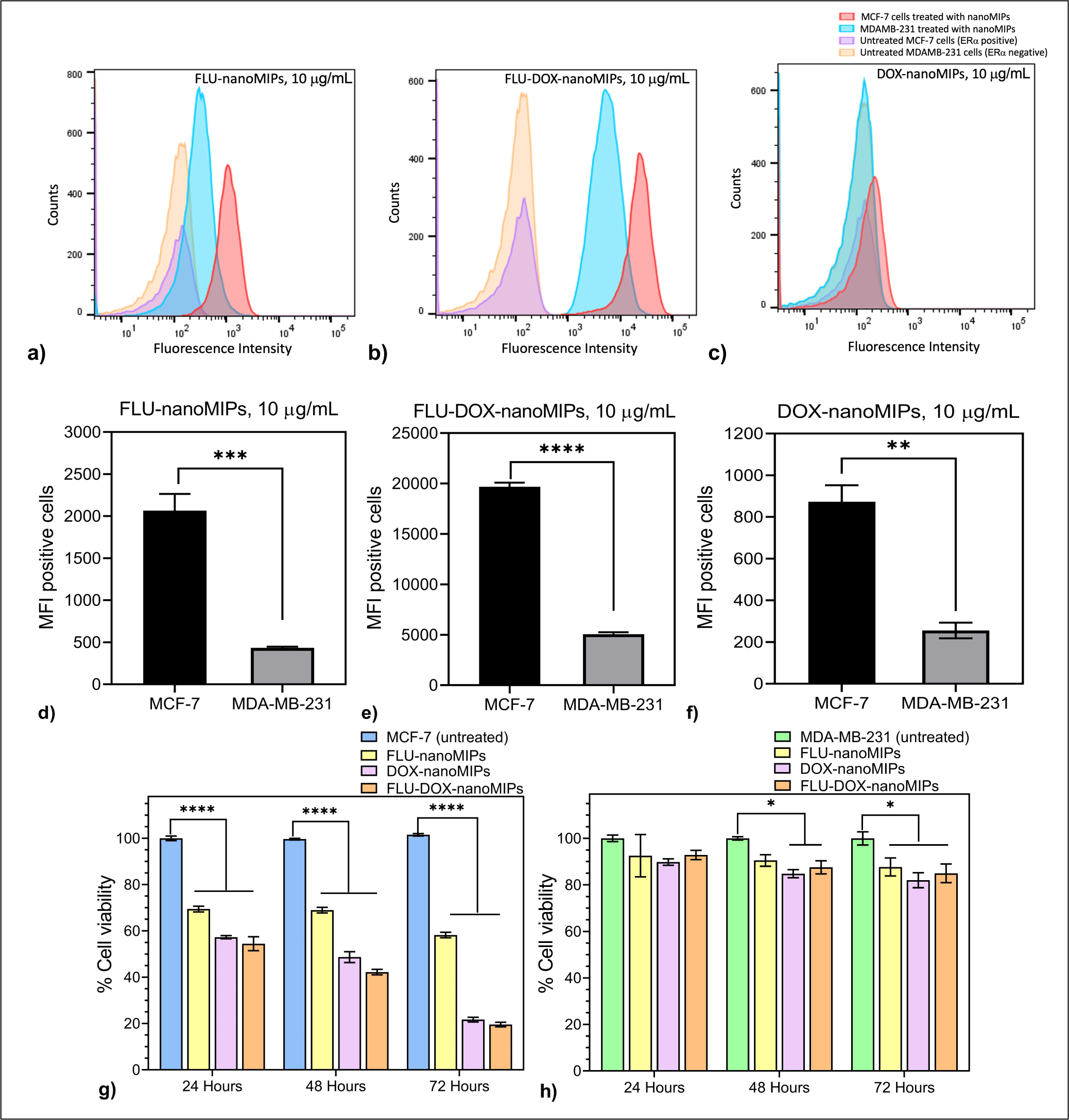
*In-vitro* flow cytometry binding assay, cells were incubated with 10 μg/mL of **a)** FLU-nanoMIPs, **b)** FLU-DOX-nanoMIPs, **c)** FLU-DOX-nanoMIPs; Mean Fluorescein intensity (MFI) of binding to MCF-7 vs MDA-MB-231, **d)** FLU-nanoMIPs, **e)** FLU-DOX-nanoMIPs, **f)** DOX-nanoMIPs. *In-vitro* cell viability assay, at 10 μg/mL for each treatment of FLU-nanoMIPs, DOX-nanoMIPs and FLU-DOX-nanoMIPs in **g)** MCF-7 and **(h)** MDA-MB-231. Data is expressed as mean ± SEM of three measurements. ** P ≤ 0.01, *** P ≤ 0.001, **** P ≤ 0.0001 *vs.* MCF-7 **(d-f)**. **** P ≤ 0.0001 *vs.* MCF-7 **(g)**, * P ≤ 0.05 *vs.* MDA-MB-231 **(h)**.

In addition to that, binding of each nanoMIPs with MCF-7 cells at higher concentration, 40μg/mL was also tested, and the comparison graph is shown in **Figure S3, supporting information**. The binding was found to be similar for each nanoMIP type, regardless of concentration. This suggests that the binding is mostly specific, and the lower concentration (10μg/mL) of nanoMIPs was sufficient to occupy most of the ERα. The comparison between FLU-DOX-nanoMIPs and FLU-nanoMIPs revealed that the double imprinting of DOX in these nanoMIPs did not have an adverse effect on their binding performance. In fact, the binding capability was observed to be improved in the double imprinted nanoMIPs.

### 2.4 *In Vitro* Cytotoxicity Assessment

*In vitro* cytotoxicity of the different nanoMIPs has been assesed using a MTT assay. After treating MCF-7 cells with different ER-α nanoMIPs (FLU-nanoMIPs, FLU-DOX-nanoMIPs, and DOX-nanoMIPs), an increase in cell cytotoxicity was observed in a time-dependent manner after 24, 48, and 72 h of incubation. Specifically, FLU-nanoMIPs demonstrated significant cytotoxicity (30.6±1.2%) after 24 h of incubation with MCF-7 cells, with only a 11% increase in cytotoxicity (total 41.8%±1.2%) after 72 hours. Conversely, FLU-DOX-nanoMIPs and DOX-nanoMIPs exhibited significant cytotoxicity in MCF-7 cells (P ≤ 0.0001) even after 24 h (42.8±0.7% and 45.5±3.0%). Furthermore, after 72 h, treatment with FLU-DOX-nanoMIPs and DOX-nanoMIPs resulted in 80.4±0.99% and 78.3±1.03% cell death, respectively (**Figure 4g**). The cytotoxicity of FLU-nanoMIPs, FLU-DOX-nanoMIPs, and DOX-nanoMIPs towards MDA-MB-231 cells was found to be only 12.3±2.7%, 15.0±2.8%, and 18±2.2%, respectively (as illustrated in Figure 4h). The MTT assay results indicated a marked difference in cytotoxicity between MCF-7 (ERα positive) and MDA-MB-231 (ERα negative) cell lines, which could be attributed to the selective binding and cellular uptake behaviour of these nanoMIPs towards ERα positive cells. The FLU-nanoMIPs exhibited selective cytotoxicity, while FLU-DOX-nanoMIPs and DOX-nanoMIPs show enhanced cytotoxicity to MCF-7 cells. This could be due to their specific binding to the helix 12 (H12) region of the ER receptor, which plays a crucial role in dimerization and transcriptional activation. This binding may explain their cytotoxic effects on MCF-7 cells, which has been previously established in various reports.

We also examined the cytotoxicity of unloaded and DOX-loaded non-imprinted nanoparticles (FLU-NIPs and FLU-DOX-NIPs) in MCF-7 and MDA-MB-231 cells. Results showed that FLU-NIPs were highly biocompatible as both cell lines demonstrated cell viability of >90% even after 72 h of exposure. However, treatment with FLU-DOX-NIPs resulted in some cytotoxicity to MCF-7 and MDA-MB-231 cells (20±0.73% and 25±2.8% after 72 h respectively) likely due to the diffusion of DOX from the nanoparticles over time (as seen in **Figure S4 a and b**). In addition, incubation with DOX showed a significant decrease in cell viability in both cell lines, which was consistent with previous literature reports [44,45].

### 2.5 CLSM imaging

To assess the cellular uptake and internalization of nanoMIPs in BC cells, CLSM microscopy has been employed. MCF-7 and MDA-MB-231 cells (∼80% confluency) were treated with FLU-nanoMIPs and FLU-DOX-nanoMIPs (10 μg/mL) in the chamber plates for 1h and 24 h at 37 °C. Following the nanoMIPs incubation, the wells were washed with PBS (three times) to remove the nanoparticles that were not internalized or bound to the cells. DAPI and Alexa Fluor™ 594 wheat germ agglutinin (WGA) was used to stain cell nucleus (blue) and plasma membrane (red) respectively. Following 1h incubation, FLU-nanoMIPs (**Figure 5 a**) and FLU-DOX-nanoMIPs (**Figure 5 b**) were found to specifically bind with the plasma membrane of the MCF-7 cells. On the other hand, minimal binding of FLU-nanoMIPs **(Figure S5)** and FLU-DOX-nanoMIPs (**Figure 5 c**) with MDA-MB-231 cells were observed. Moreover, difference in MFI (calculated from confocal images) between MCF-7 cells and MDA-MB-231 cells treated with FLU-nanoMIPs (P ≤ 0.01) and FLU-DOX-nanoMIPs (P ≤ 0.001) was found to be statistically significant (**Figure 5 d**). This signifies that the binding of nanoMIPs is specific to the ERα positive cells.

**Figure 5:**
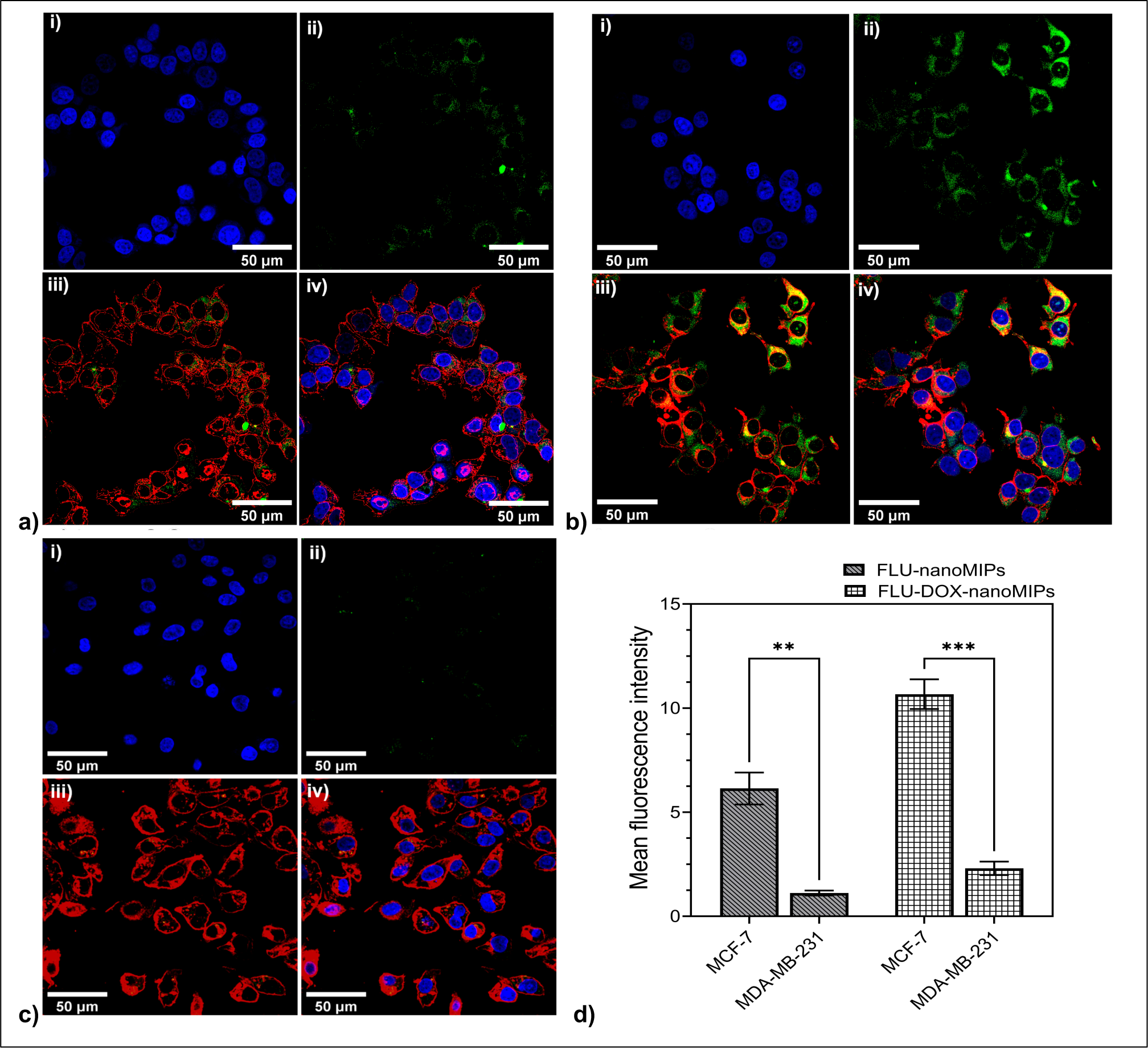
CLSM images (40X) of MCF-7 cell line incubated for 1 h (at 37 °C) with **(a)** FLU-nanoMIPs and **(b)** FLU-DOX-nanoMIPs; **(c)** CLSM images (40X) for MDA-MB-231 incubated with FLU-DOX-nanoMIPs for 1 h at 37 °C (i) DAPI, (ii) FLU-DOX-nanoMIPs with green fluorescence (iii), plasma membrane with red fluorescence (WGA antibody Alexa Fluor™ 594) with FLU-DOX-nanoMIPs (iv), merged **(d)** The mean fluorescence intensity of FLU-nanoMIPs and FLU-DOX-nanoMIPs in MCF-7 cells and MDA-MB-231 cells. ** P ≤ 0.01, *** P ≤ 0.001, *vs.* MCF-7.

After 24h, it was evident that FLU-nanoMIPs and FLU-DOX-nanoMIPs (green) were able to cross the plasma membrane and get internalized in to MCF-7 cells (**Figure 6 a and b**). On the contrary, no significant uptake of nanoMIPs in the ERα negative cell line MDA-MB-231 was observed, as shown in **Figure 6 c and d**. It has been reported that ERα binds with Caveolin-1, which is one of the structural proteins of caveolae, and plays a major role in trafficking of ERα to and from the cell surface and maintaining an environment for cell signalling and endocytosis of ERα [46].Thus, the internalization of nanoMIPs into MCF-7 cells can likely be attributed to the ERα mediated caveolae dependent endocytosis [8,47]. Furthermore, the difference in MFI between MCF-7 cells and MDA-MB-231 cells subjected to 10 μg/mL of FLU-nanoMIPs (P ≤ 0.01) and FLU-DOX-nanoMIPs (P ≤ 0.001) was statistically significant **(Figure 6 e).**

**Figure 6:**
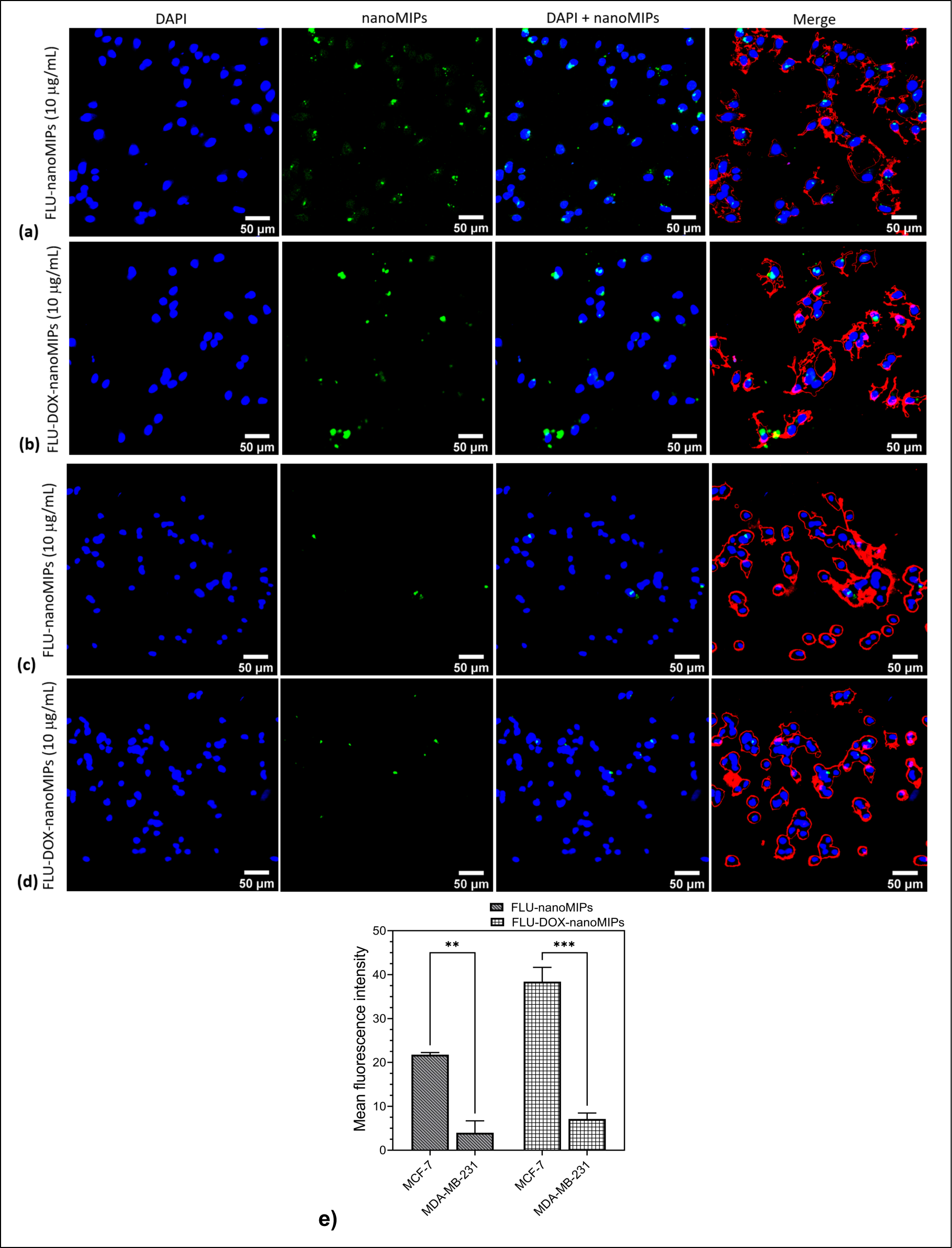
CLSM images (20X) for nanoMIPs incubated for 24 hours at 37 °C **(a)** MCF-7 cells incubated with FLU-nanoMIPs **(b)** MCF-7 cells incubated with FLU-DOX-nanoMIPs **(c)** MDA-MB-231 cells incubated with FLU-nanoMIPs **(d)** MDA-MB-231 cells incubated with FLU-DOX-nanoMIPs; Nucleus is stained with blue fluorescence (DAPI), nanoMIPs with green fluorescence, membrane with red fluorescence (WGA antibody Alexa Fluor™ 594) **(e)** The mean fluorescence intensity of FLU-nanoMIPs and FLU-DOX-nanoMIPs in MCF-7 cells and MDA-MB-231 cells. ** P ≤ 0.01, *** P ≤ 0.001, *vs.* MCF-7.

After internalization (as illustrated at 63X in **Figure 7 a** for FLU-nanoMIPs and **Figure 7 b** for FLU-DOX-nanoMIPs), the nanoMIPs proceed to translocate into the nucleus, as demonstrated in **Figure 7 ci**. The 3D CLSM image confirmed the colocalization of the nanoMIPs within the nucleus of ERα positive cells, as evidenced by the overlapping fluorescence signal in yellow (**Figure 7 cii**). It has been observed that 3.53% of the volume of the nucleus (in case of **Figure 7 cii**) colocalizes with FLU-DOX-nanoMIPs, furthermore, 15.61% and 1.71% of the volume of Cluster 1 and Cluster 2 respectively colocalized with the nucleus.

**Figure 7:**
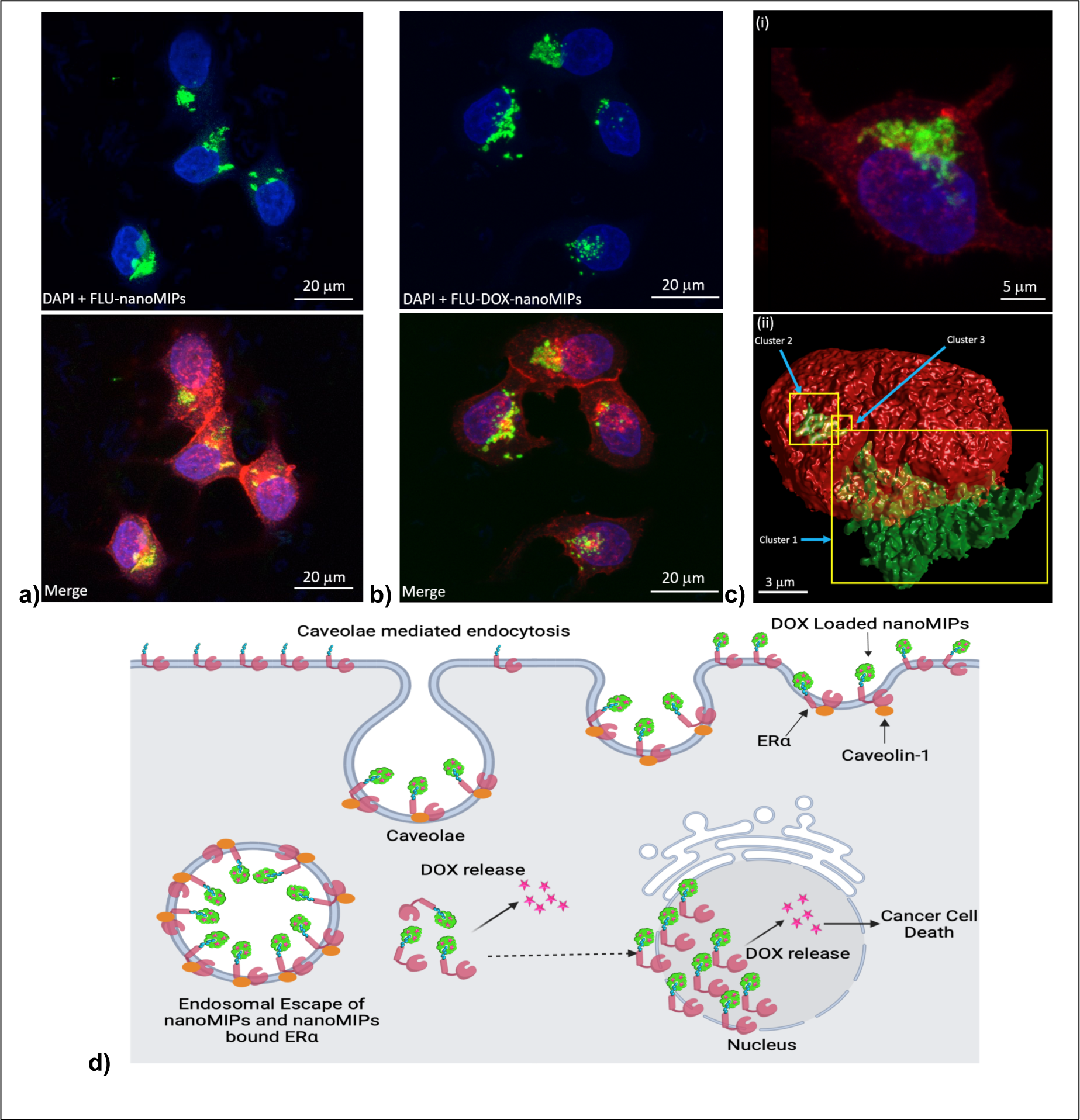
CLSM images (63X) of MCF-7 cell line incubated with **(a)** FLU-nanoMIPs, **(b)** FLU-DOX-nanoMIPs for 24 hours at 37 °C, **(c)** 3D image of FLU-DOX-nanoMIPs internalization in MCF-7 cells **(i), 3D render image** showing the yellow areas (indicated by the blue arrows) are FLU-DOX-nanoMIPs (green) clusters colocalizing (hence, inside) the nucleus (red) of ERα positive MCF-7 cells **(ii).** Nucleus is stained with blue fluorescence (DAPI), nanoMIPs with green fluorescence, membrane with red fluorescence (WGA antibody Alexa Fluor™ 594). **(d)** Schematic representation of internalization of nanoMIPs through caveolae mediated endocytosis membrane ERα,) followed by translocation to the nucleus and release of DOX in cytoplasm as well as in nucleus.

The ability of DOX loaded nanoMIPs to specifically bind to ERα positive MCF-7 and its nucleus led to the release of DOX to both cytoplasm and nucleus, thus resulting in higher cytotoxicity compared to MDA-MB-231 cells treated with nanoMIPs. These findings were consistent with our cytotoxicity results observed in MTT assay even at a lower concentration of DOX (∼2 μg) in presence of 10 μg/mL nanoMIPs.

### 2.6 Evaluation of the action of nanoMIPs against breast cancer cells in a 3D biomaterial-based model

In order to investigate the action of nanoMIPs against MCF-7 and MDA-MB-231 cells, in a more tissue biomimetic environment, we conducted experiments in 3D porous polymer (PU) scaffolds, surface modified with Collagen I, for extracellular matrix (ECM) mimicry (**Figure 8a**) [48,49,50]. Collagen I is known to influence various factors related to the tumour microenvironment in breast cancer, such as proliferation, survival, migration, and invasion. Changes in the structure of Collagen I have been observed during early breast cancer development and are associated with local invasion. Additionally, the density of Collagen I has been found to correlate with the progression of breast cancer [51,52] Overall, the 3D scaffolds are a better mimicry of essential aspects of tumour microenvironment (*in vivo*) in terms of structure, cell-to-cell and cell-to-extracellular matrix interactions and spatial orientation, as compared to the simple 2D culture shown in the previous sections [48,49].

**Figure 8:**
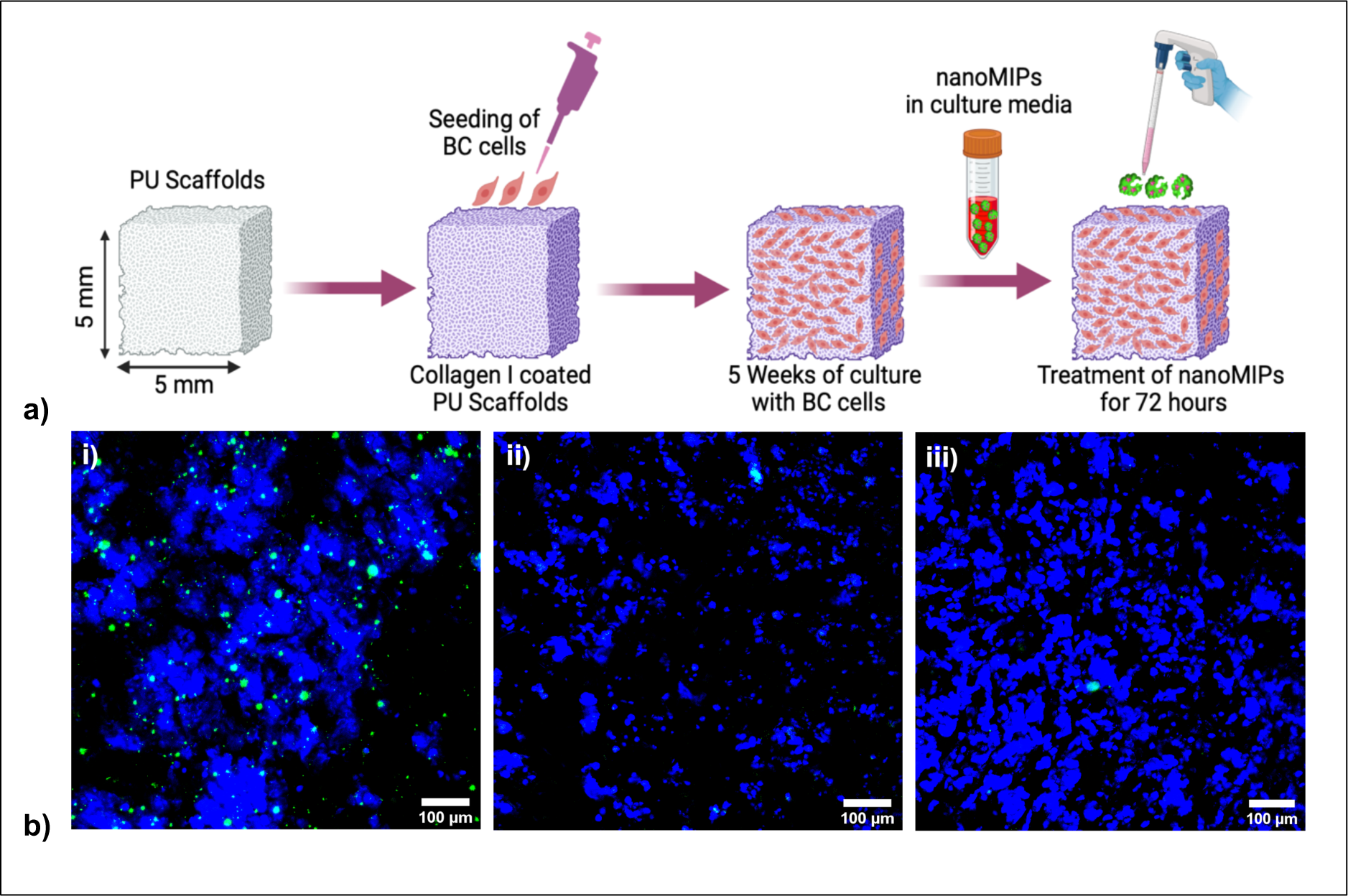
**(a)** Schematic of cell culture seeding and treatment of nanoMIPs to 3D scaffold; **(b)** CLSM images (10X) of 3D scaffolds (poly-urethane scaffolds coated with Collagen-1) of MCF-7 cell lines treated with **i)** FLU-nanoMIPs, **ii)** FLU-DOX-nanoMIPs, and **iii)** MDA-MB-231 3D scaffolds treated with FLU-DOX-nanoMIPs (after 72 h treatment) at 37 °C. Nucleus is stained with DAPI (Blue); nanoMIPs are in green.

More specifically, as described in the methodology section, at week 5 of the 3D culture, the scaffolds were exposed to various treatments for 72 hours after which analysis of their spatial distribution, viability and apoptosis took place (please see section 4.13).

Our results showed that FLU-nanoMIPs successfully penetrated the 3D scaffolds of ERα-positive MCF-7 cell lines, as indicated by the green fluorescence in **Figure 8 bi**. Clear binding of FLU-nanoMIPs with the nucleus has also been observed (co-localization). These results evidenced that these nanoMIPs can penetrate into complex 3D *in vitro* models.

In the case of FLU-DOX-nanoMIPs (**Figure 8 bii**), binding to the 3D scaffolds of ERα-positive MCF-7 cell lines was observed, but a smaller number of cells and lower fluorescence intensity for nanoMIPs were detected compared to FLU-nanoMIPs. This suggests that after treatment with DOX-loaded nanoMIPs, the cells that have up-taken these nanoMIPs underwent cell death, detached from the scaffolds and were eventually washed away during the washing process.

Conversely, minimal binding of FLU-DOX-nanoMIPs (green fluorescence), and appreciable cell density was observed in the 3D scaffolds of ERα-negative MDA-MB-231 cell lines (**Figure 8 biii**). These results confirm the specific targeting ability of nanoMIPs towards breast cancer cells within the complex network of the 3D scaffolds. The lack of specific uptake in MDA-MB-231 cells further supports our findings from the 2D cell line experiments.

#### Conclusions

In conclusion, the development of doubly-molecularly imprinted polymeric nanoparticles (nanoMIPs) offers a promising tool for targeted drug delivery in precision medicine BC treatment. The synthetic approach used to produce such nanoMIPs relied on a modified solid-phase methodology, which enables the formation of two types of binding sites: one specific towards the ER, the other to a drug (DOX), which enables the use of these nanoparticles as drug delivery systems. Synthesized DOX loaded nanoMIPs with spherical morphology typical size ranging from 140-170 nm. These materials rival the affinity of commercial antibodies, with a K_D_ of 10 nM for ERα receptor as determined by SPR measurements. These double imprinted nanoMIPs demonstrated to target both membrane and nuclear ERα, thus enabling nuclear DOX delivery. Moreover, these nanoMIPs specifically bind and elicit cytotoxicity (∼80%) to the ERα positive cancer cells compared to ERα negative cell lines (∼15%). This suggests that these smart nanosystems, if used as drug delivery agents, can potentially minimize the off-target side-effects while improving on drug efficacy. It is worth noting that the membrane to nuclear drug delivery behaviour observed in this work is entirely novel and has not been reported on with molecularly imprinted or other hybrid nanoparticle systems. Importantly, ERα is also highly expressed in other types of cancer, such as endometrial, prostate, and ovarian cancer. Given the versatility of nanoMIPs, it is straightforward to extend this technology to the treatment of other cancers or other drug compounds. However, further studies are needed to explore the potential of these nanoparticles in clinical settings, their *in viv*o biodistribution and biocompatibility, and optimize their efficacy and safety. Overall, the use of targeted drug delivery systems such as nanoMIPs hold great promise for the future of cancer treatment due to their cost-effective nature, robustness, high selectivity, and scalable production process.

## 4.0 Materials and methods

### 4.1 Materials

Glass beads (53-106 μm diameter, Spheriglass 2429 CP00) were purchased from Blagden Chemicals (Kent, United Kingdom). *N*-Isopropylacrylamide (NIPAM), *N,N′-* Methylenebis(acrylamide) (Bis), *N*-*tert*-Butylacrylamide (nTBA), *N-*(3-Aminopropyl)methacrylamide hydrochloride (APMA), acrylic acid (AA), Fluorescein O-methacrylate, N,N,N’,N’-tetraethylethylenediamine (TEMED), (3-aminopropyl)trimethoxy-silane (APTMS), doxorubicin HCl (DOX), dialysis cartridges (Vivaspin^®^ 20, 3 kDa molecular weight cut-off (MWCO) Polyether-sulfone), Supelco polypropylene solid phase extraction tubes (60 mL) and 3-[4, 5-dimethylthiazol-2-yl]-2,5-diphenyltetrazolium bromide (MTT), 1-Ethyl-3-(3-dimethylaminopropyl) carbodiimide (EDC), dipotassium phosphate, disodium phosphate, ethanolamine, *N*-hydroxysuccinimide (NHS), and Tween 20 were purchased from Sigma Aldrich, Poole, Dorset, UK. PierceTM Bicinchoninic Acid Assay (BCA) protein assay kit, ammonium persulfate (APS), methanol, acetone, acetonitrile, sodium hydroxide (NaOH), hydrochloric acid (33%, HCl), succinimidyl iodoacetate (SIA), potassium chloride, sodium acetate, sodium chloride, Oxoid phosphate buffered saline (PBS) tablets (Catalog number: BR0014G) and human recombinant ERα protein (Accession number NP_000116.2, Catalog number: A15674), Alexa Fluor™ 594 Wheat Germ Agglutinin (WGA) antibody, Caspase-3/7 green detection reagent, a Live/Dead Viability/Cytotoxicity Kit (Molecular Probes) and DAPI were purchased from Fisher Scientific UK Ltd (Loughborough, UK). CSHSLQKYYITGEAEGFPATV epitope of ERα was synthesized by Elabscience^®^ and obtained through Caltag Medsystems (Buckingham, UK). All chemicals and solvents were high-performance liquid chromatography (HPLC) analytical grade and were used without any further purification. PBS solutions were prepared with deionized (DI) water with resistivity of ≥18.2 MΩ cm.

### 4.2 Preparation of Epitope derivatised glass beads

60 g of glass beads (53-106 μm) were activated by boiling in 2 M NaOH (24 mL) for 15 min, then washed with double-distilled water (10 times with 100 mL) until the pH of the washed solution was around 7.4. Afterwards, the glass beads were rinsed twice with acetone (100 mL) and dried at 80°C for 2 h. Subsequently, the activated glass beads were incubated in a 24 mL solution of 2%, v/v APTMS in anhydrous toluene for 12 h for the silanization step. After the incubation step, the glass beads were transferred into sintered funnel and washed with acetone (8 × 50 mL) and methanol (3 × 50 mL), subsequently dried under vacuum. Then, 60 g of glass beads were placed in a solution of SIA (0.2 mg/mL in acetonitrile) for 2 h in the dark (0.4 mL solution/g glass beads). Afterwards, the beads were washed with 400 mL of acetonitrile in a sintered glass funnel and incubated with 7 mg of cysteine-modified peptide epitope (primary template) in 40 mL of deoxygenated 1× phosphate buffered saline (PBS) containing 5 mM EDTA, pH 8.2. Covalent immobilisation of the epitope on the silanized bead was confirmed by performing a BCA assay [53].

### 4.3 Synthesis of nanoMIPs/NIPs against ERα

The synthesis protocol was adapted from previous reports of Canfarotta and colleagues [54]. A monomer mixture containing NIPAM (39 mg), Bis (2 mg), nTBA (33 mg, pre-dissolved in ethanol 1 mL), APMA (5.8 mg), AA (2.2 μL), and Fluorescein O-methacrylate (FLU, 2.6 mg) were dissolved in 100 mL of PBS (5 mM, pH = 7.4). To fabricate DOX loaded doubly molecularly imprinted nanoparticles (DOX-nanoMIPs) and fluorescein tagged DOX-nanoMIPs (FLU-DOX-nanoMIPs), 3 mg of DOX was added to the monomer mixture. Fluorescein tagged nanoMIPs without DOX (unloaded) are named as FLU-nanoMIPs. Non-imprinted nanoparticles (NIPs) were produced according to the aforementioned method, except for the substitution of epitope-derivatized beads with silanized beads. Briefly, the solution containing the monomer mixture was degassed under vacuum and sonicated for 10 min, and purged with N_2_ for 30 min.

Then, 60 g of ERα epitope derivatized glass beads were added to the solution under continuous N_2_ purging, and polymerization was initiated by adding mixture containing 800μL of APS aqueous solution (60 mg/mL) and 24 μL of TEMED. Mixture was kept for 4 hours at room temperature (RT) for polymerization, afterwards, poured into a solid-phase extraction (SPE) cartridge (60 mL) fitted with a frit (20 μm porosity). The removal of low affinity nanoMIPs, polymer and unreacted monomers was achieved by washing with distilled water (9 × 20 mL) at RT. Following that, 20 mL of distilled water pre-warmed at 65°C was poured into the SPE and was placed in a water bath at 65°C for 15 min. This step was repeated 5 times, until ∼100 mL of high-affinity nanoMIPs solution were collected. Concentrated nanoMIPs were obtained by evaporating the dispersion solvent in an oven for 24 hours at 60°C. The obtained nanoMIPs were further purified using a centrifugal dialysis cartridge fitted with a membrane of 3 kDa MWCO. Five washes with deionized water (10 mL) were performed n obtained nanoMIPs were re-suspended in 50 mL deionized water.

### 4.4 Determination of drug loading from DOX-loaded nanoMIPs

A stock solution of DOX (100 μg/mL) was prepared in water and further diluted in deionized water to the concentration of 1, 2.5, 5, 7.5 and 10 μg/mL. UV-visible spectra were recorded between 200 to 600 nm using a Jenway 7200 UV-Visible scanning spectrophotometer. The DOX calibration curve was plotted using absorbance (α_max_ 254 nm) *vs.* concentration (**Figure S1**) and linear calibration curve equation (***y=mx+b)*** was used to estimate the DOX within the nanoMIPs. The calibration curve was plotted in the UV region at 254 nm because in the visible region fluorescein-o-methacrylate peak overlap with the visible region peaks of DOX. The percent drug loading capacity (DLC) was calculated by taking the ratio of amount of DOX encapsulated and weight of nanoMIPs

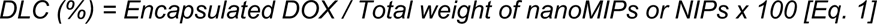

Drug loading efficiency of DOX loaded into nanoMIPs have been calculated with following formula.

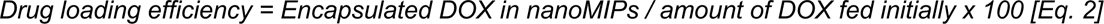

### 4.5 Dynamic light scattering (DLS) and Electrophoretic Light Scattering measurements

Dynamic light scattering (DLS) experiment were performed using Malvern Zetasizer Nano ZS to measure the hydrodynamic diameter (*D_h_*) of different nanoMIPs at 25±0.1°C. The instrument used scattering angle of 173° and the laser wavelength of 632.8 nm. The size was measured at different times to evaluate the stability of the systems at 25 ± 0.1°C. The dispersion of nanoMIPs in distilled water was subjected to sonication for 30 minutes and subsequently examined by DLS inside a 3 mL disposable polystyrene cuvette. Stability of nanoMIPs was determined with same instrument employed for DLS measurements.

### 4.6 Morphology of nanoMIPs

For transmission electron microscopy (TEM), a nanoMIP solution (40 μg/mL) was drop-casted on the copper grids and images was captured using Hitachi HT7800 120kV TEM machine (Tokyo, Japan) equipped with EMSIS CMOS Xarosa camera. For scanning electron microscopy (SEM) analysis, nanoMIP solutions were drop casted and dried on the glass chips (1×1 cm) and measurements were performed using a Tescan Vega 3LMU (Kohoutovice Czech Republic) machine with tungsten filament.

### 4.7 Immobilisation of nanoMIPs onto the SPR Sensor Surface

Carboxymethyl dextran hydrogel coated Au chips, purchased from Reichert Technologies (Buffalo, USA) were installed onto a Reichert 2 SPR following the manufacturer’s instructions. The sensor surface was then preconditioned by running buffer PBST (PBS pH 7.4 and 0.01 % Tween 20) at 10 µL min^-1^ until a stable baseline was obtained. The flow rate of 10 µL was maintained throughout the immobilisation process. To activate carboxyl groups on the surface of the sensor chip, a freshly prepared aqueous solution (1 mL) of EDC (40 mg) and NHS (10 mg) was injected onto the sensor chip surface for 6 minutes.

To the activated surface, 300 µg of the nanoMIPs dissolved in 1 mL of the running buffer (PBST) and 10 mM sodium acetate were injected only to the left channel of the surface for 1 minute. Finally, a quenching solution (1 M ethanolamine, pH 8.5) was injected for 8 min to deactivate carboxyl groups and wash away the unbound nanoMIPs. A continuous flow of running buffer (PBST) at 10 µL min-1 was maintained after nanoMIPs immobilisation. SPR measurements were carried out after a stable baseline was achieved. The left channel was the working channel, and the right channel was the reference.

### 4.8 Kinetic Analysis Using SPR

Kinetic analysis was initiated by injection of the running buffer PBST (blank) onto the nanoMIPs immobilised sensor surface for 2 min, followed by PBST for 5 min. The binding kinetics of an individual nanoMIPs to the selected target was determined from serial dilutions (five concentrations in the 4-64 nM range) of the selected target under study. Each dilution was injected for 2 min (association) followed by PBST for 5 min (dissociation). After dissociation, the target was removed from the immobilized surface by injecting regeneration buffer (10 mM Glycine-HCl, pH 2) for 1 min followed by PBST for 1 min. The same procedures were repeated for the remaining four dilutions of the target. After the analyses were completed, signals from the left channel were subtracted from signals from their respective reference channel (the right channel).

The SPR responses from five concentrations of the target compound (4-64 nM) were fitted to a 1:1 bio-interaction (BI) model (Langmuir fit model) utilizing TraceDrawer software. Association rate constants (*k_on_*), dissociation rate constants (*k_off_*), and maximum binding (*B_max_*) were fitted globally, whereas the BI signal was fitted locally. The equilibrium dissociation constant (*K_D_*) was calculated from the ratio *k_off_/k_on_*.

### 4.9 Cell culture conditions

MCF-7 and MDA-MB 231 (was a gift from Dr Annette Meeson, Newcastle University) were cultured in DMEM (GIBCO) supplemented with 10% foetal bovine serum (FBS, GIBCO) and 1% penicillin/streptomycin (P/S). The cells were incubated in a humidified atmosphere of 5% CO_2_ and 95% air at 37°C.

### 4.10 *In-vitro* cell binding by flow cytometry

*In-vitro* cell binding of different nanoMIPs with cancer cells has been determined using a BD LSRFortessa™ X-20 flow cytometer. Cells were gently scraped, after washing with PBS, and resuspended in flow cytometry buffer (1xPBS, 0.5% BSA, 0.1% NaN_3_) at concentration 1×10^6^ cells in 100 µL. Prior to treatment, nanoMIPs were sonicated (10 min) and added to the cell suspension with the final concentration of 10 µg mL^-1^ and 40 µg mL^-1^. Following the 2 h incubation, the cells were centrifuged, washed, and dispersed in flow cytometry buffer and subsequently, binding analysis was performed using flow cytometer. Each experiment was done in triplicate. Data was analyzed using flowJo software, version 10.

### 4.11 *In-vitro* cytotoxicity assessment

The MTT assay was used to evaluate the *in-vitro* cytotoxicity of various nanoMIPs and pure DOX in MCF-7 and MDA-MB-231 cells [55]. For each experiment, 5000 cells/well were seeded into 96-well plates and incubated for 24 h. The cells were then treated with the synthesized nanoMIPs and pure DOX at a concentration of 10 µg/mL for 24, 48, and 72 h. 2.5 mg of MTT was dissolved in 500 µL of phosphate-buffered saline (PBS) and diluted to 5 mL with serum free DMEM medium. 200 μL of the MTT solution was added to each well of the plates after 24 h, 48h and 72h of nanoMIPs treatment respectively and the plate wrapped in aluminium foil were incubated at 37 °C for 4 h. After that, the media containing unbound MTT and dead cells was removed from each well, and 200 µL of DMSO was added to each well. The plates were shaken on orbital shaker at 70 rpm for 30 min, and the absorbance was measured at dual wavelengths of 550 nm and 630 nm using a Spectramax M5^e^ (Moecular Devices, CA) multi-mode automated microplate reader. The results were expressed as percentage cell viability, assuming the viability of control cells as 100%. Three independent experiments were performed for each study, and all measurements were performed in triplicate.

### 4.12 Confocal laser scanning microscopy (CLSM) imaging

nanoMIPs binding as well as cellular uptake in MCF-7 and MDA-MB-231 cell lines was assessed at different time points (1 and 24 h). The cells were seeded at 20000 cells/well (300 μL) in µ-Slide 8 Well high ibiTreat chamber slides (Thistle Scientific, Uddingston, Glasgow, UK) and kept for 24 h to achieve 70-80% confluency. Prior to the treatment, nanoMIPs were washed with ethanol (70%) and then with PBS (two times) before adding to the cells, using centrifugation cartridges.

After 24 h, the medium was pipetted out and the nanoMIPs suspension (10 µg/mL, in full media) was added to the chamber plates; then the plates were incubated for 1 h or 24 h. Following the treatment, the chamber plates were washed three times with PBS (pre warmed) to remove the unbounded nanoMIPs. Afterwards, the treated cells were fixed using 4% PFA solution prepared in PBS for 20 min at room temperature. PFA fixed chamber wells were then incubated with wheat germ agglutinin (WGA) with Alexa Fluor™ 594 (catalogue no. W11262) for 10 minutes (dilution 1:200) to stain plasma membrane. After washing the chamber wells with PBS (3 times, 10 min each), DAPI (1:200 in PBS) was added for 20 minutes. The chamber plates were washed with PBS (three times) and then images were taken with a confocal laser scanning microscope Leica TCS SP8 STED 3X (Leica microsystems, Germany) using the following lasers (i) 405nm for DAPI (blue), (ii) 488nm for fluorescein tagged nanoMIPs (green) and (iii) 561nm for Alexa Fluor 594 (red). The images were analyzed using Image J® software (Wayne Rasband, NIH, Bethesda, MD, USA).

### 4.13 3D Scaffold preparation and cell culture

The thermal-induced phase separation method was utilized to synthesize 3D porous polyurethane (PU)-based scaffolds, which were then sterilized, and surface modified (coated) with Collagen I, as per the methodology described in [56,57,58].

The MCF-7 and MDA-MB-231 cell lines were seeded onto scaffolds measuring 5 × 5 × 5 mm³ at a seeding density of 0.5 × 10⁶ cells per scaffold. The cells were cultured for a duration of 5 weeks. Subsequently, incubated with various treatments, including FLU-nanoMIPs, FLU-DOX-nanoMIPs and DOX (10 μg/mL) lasting 72 hours, and the treatment was removed. After treatment, the scaffolds were characterized through sectioning, staining, and image analysis via CLSM.

## Acknowledgments

The research and innovation program, “European Union’s Horizon 2020,” has provided support for this work in the form of a Marie Sklodowska-Curie Postdoctoral fellowship awarded to Dr Pankaj Singla (grant agreement number-893371, TEMPER). 3D cell line work in this manuscript is possible from the funding of Newcastle Wellcome Trust Translational Partnership (NWTTP). Dr Eirini Velliou is grateful to the Medical Research Council UK for a New Investigator Research Grant (MR/V028553/1), which also financially supports Dr Priyanka Gupta. Authors would like to thank Dr Annette Meeson, International Centre for Life, Newcastle University for generously proving the breast cancer cell lines and valuable discussion.

