## Supplementary material for "Double Imprinted Nanoparticles for Sequential Membrane-to-Nuclear Drug Delivery": Supplementary Information.docx

| Batch | ERa epitope | NIPAM | nTBA | AA | APMA | Bis | FLU | DOX |
| --- | --- | --- | --- | --- | --- | --- | --- | --- |
| FLU-nanoMIPs | + | + | + | + | + | + | + | - |
| FLU-DOX-nanoMIPs | + | + | + | + | + | + | + | + |
| DOX-nanoMIPs | + | + | + | + | + | + | - | + |
| FLU-NIPs | - | + | + | + | + | + | + | - |
| FLU-DOX-NIPs | - | + | + | + | + | + | + | + |

**Table S1:** Composition of different batches of the nanoMIPs and NIPs fabricated in this study.

**Table S2:** DOX loading, loading efficiency and loading capacity of different batches of DOX loaded nanoMIPs and NIPs.

|  | Loading DOX concentration (μg/100 μg) | Loading efficiency | Loading Capacity |
| --- | --- | --- | --- |
| DOX-nanoMIPs | 17.28 ± 0.1 | 57.6 ± 0.33 % | 17.28 ± 0.1 % |
| FLU-DOX-nanoMIPs | 19.33 ± 0.16 | 64.43 ± 0.53 %, | 19.27 ± 0.16 % |
| FLU-DOX-NIPs | 18.37 ± 0.12 | 61.12 ± 0.40 %. | 18.21 ± 0.12% |

**a)**
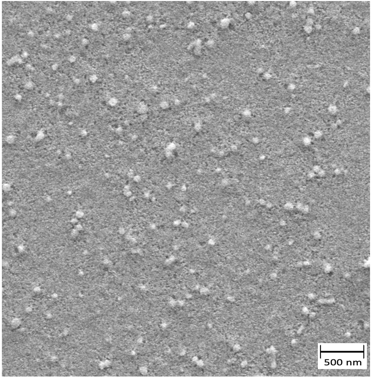
 **b)**
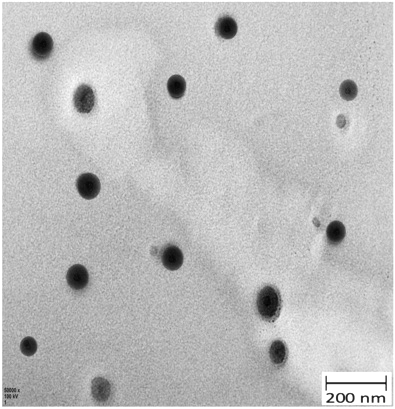

**Figure S1:** Characterization: **a)** representative SEM images of DOX-nanoMIPs; **b)** TEM image (25000x) DOX-nanoMIPs

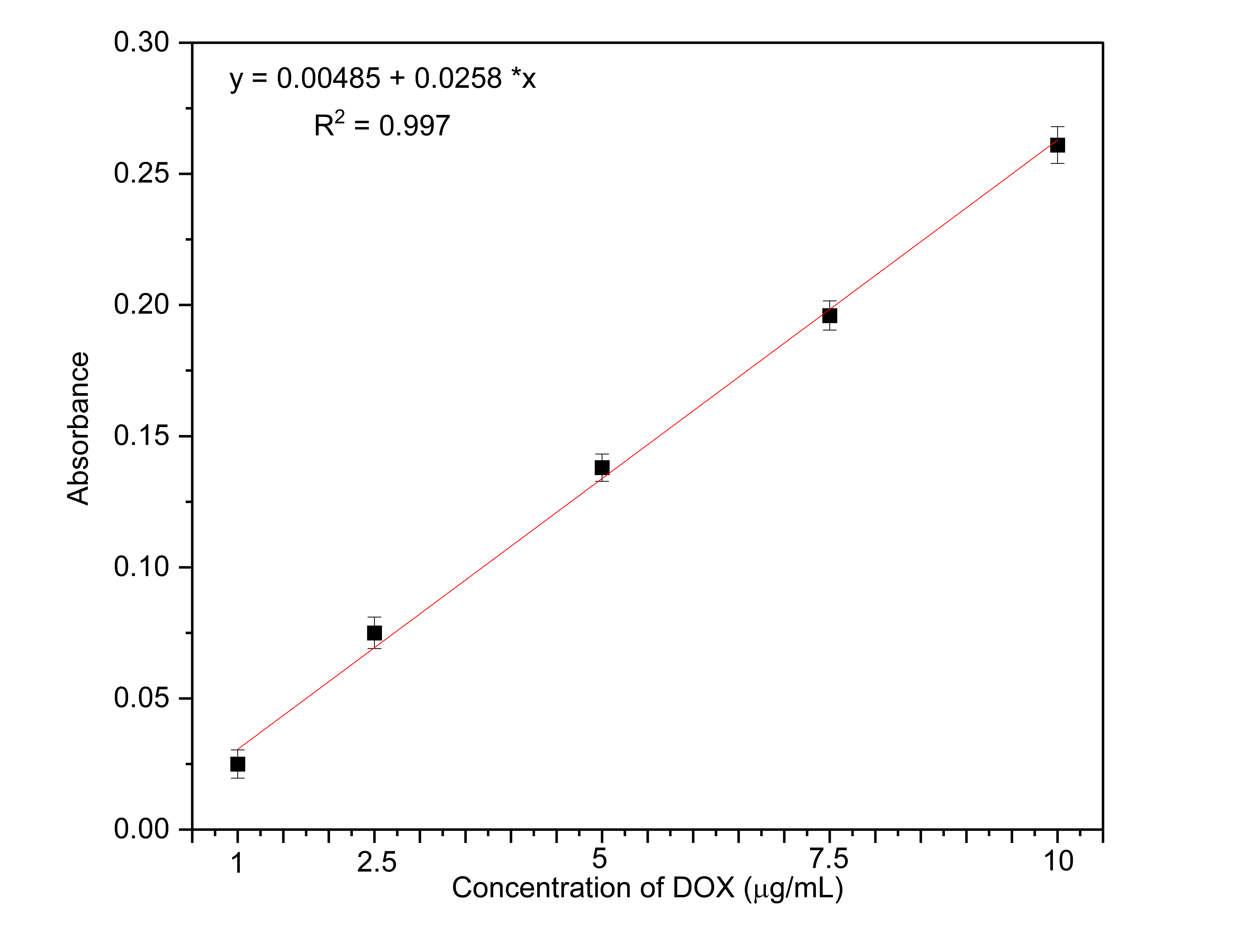

**Figure S2:** Calibration curve of DOX absorbance (l_max_ 254 nm) *vs.* concentration (1, 2.5, 5, 7.5 and 10 μg/mL) obtained from UV-visible spectras.

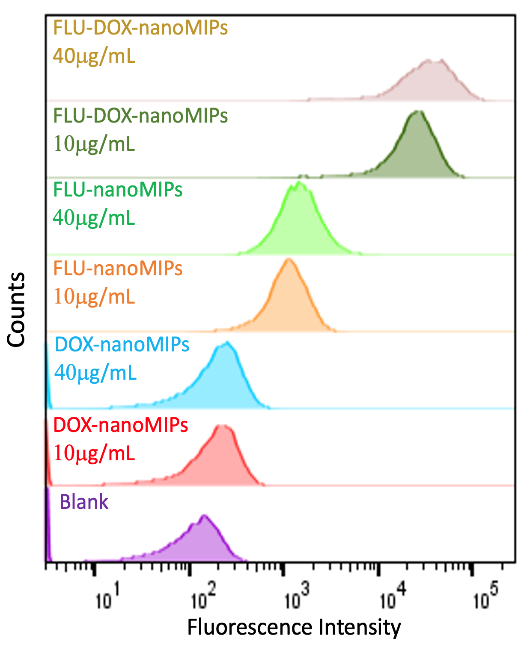

**Figure S3:** *In-vitro* flow cytometry binding assay, MCF-7 cells were incubated with 10 μg/mL and 40 μg/mL of DOX-nanoMIPs, FLU-nanoMIPs and FLU-DOX-nanoMIPs.

**a)**
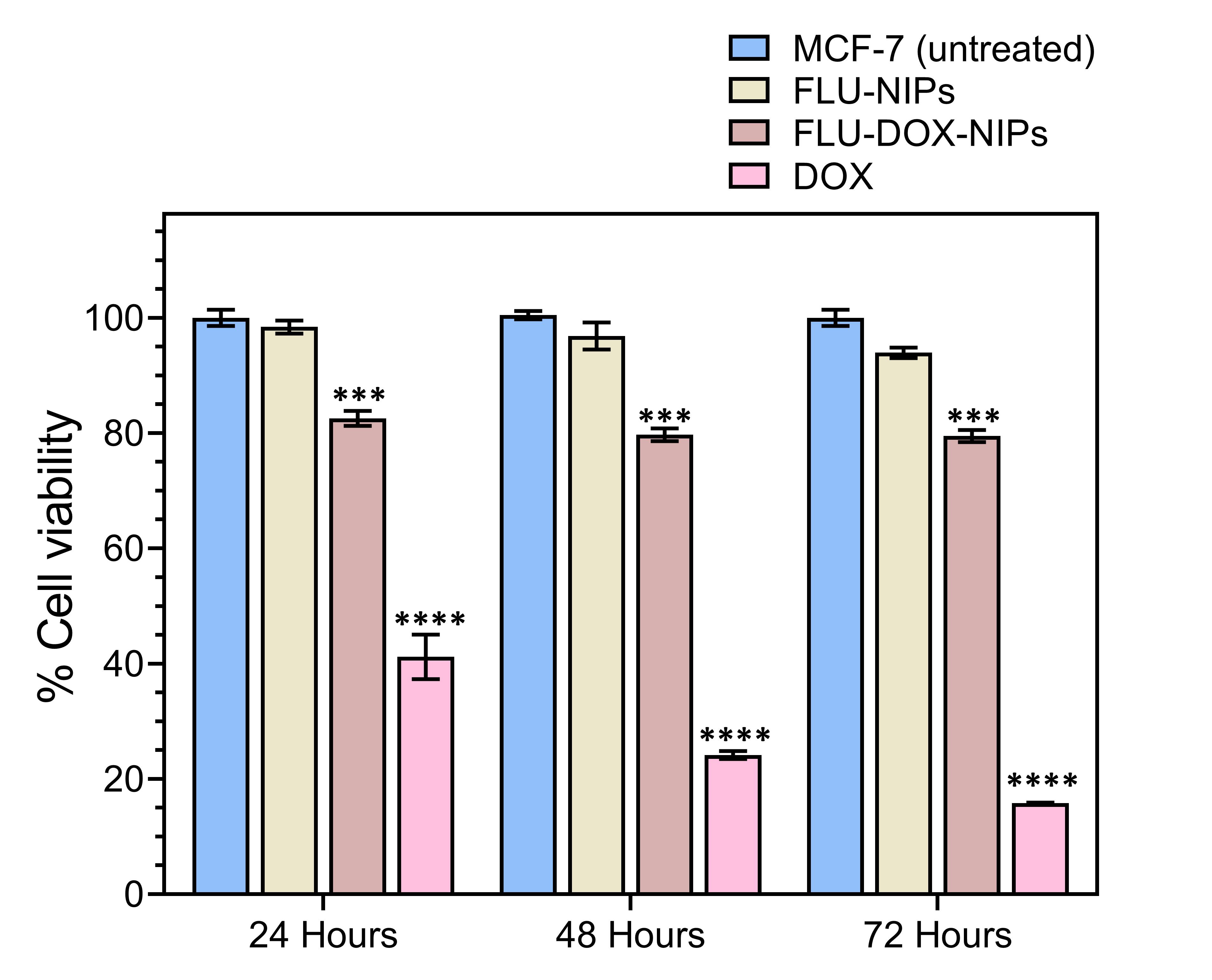
 **b)**
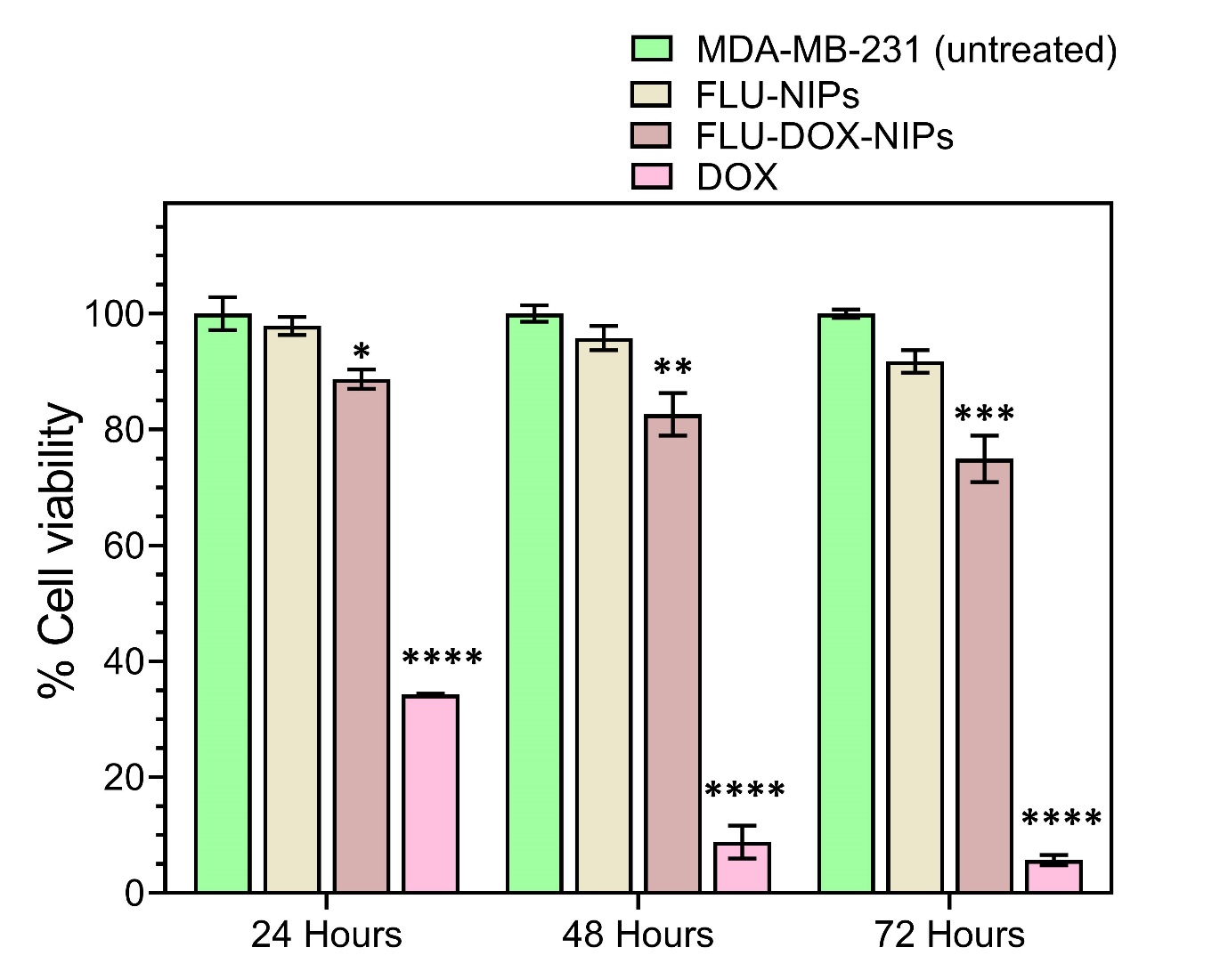

**Figure S4:** *In-vitro* cell viability assay, at 10 μg/mL for each treatment with FLU-NIPs, FLU-DOX-NIPs and DOX, **a)** MCF-7 and **b)** MDA-MB-231. Data is expressed as mean ± SEM of three measurements. *** P ≤ 0.001, **** P ≤ 0.0001 *vs.* MCF-7 control (**a**), *P ≤ 0.05, ** P ≤ 0.01, *** P ≤ 0.001, **** P ≤ 0.0001 *vs.* MDA-MB-231 control (**b**).

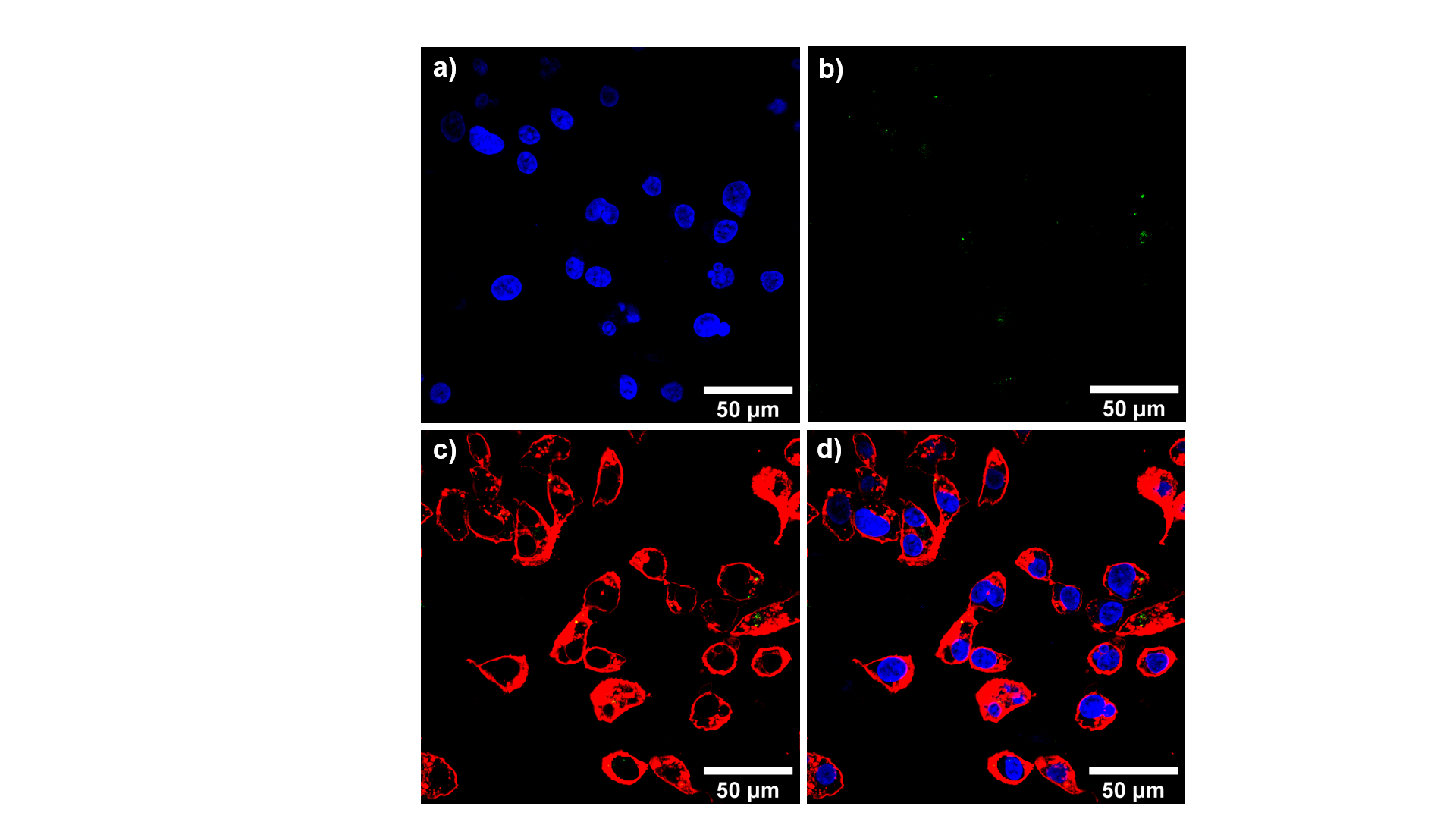

**Figure S5:** CLSM images (40X) for MDA-MB-231 incubated with FLU-DOX-nanoMIPs for 1 h at 37 °C **(a)** DAPI, **(b)** FLU-DOX-nanoMIPs with green fluorescence, **(c)** plasma membrane with red fluorescence (WGA antibody Alexa Fluor™ 594) with FLU-DOX-nanoMIPs, **(d)** merged.
